# Organelle genome assembly uncovers the dynamic genome reorganization and cytoplasmic male sterility associated genes in tomato

**DOI:** 10.1101/2021.03.03.433741

**Authors:** Kosuke Kuwabara, Issei Harada, Yuma Matsuzawa, Tohru Ariizumi, Kenta Shirasawa

## Abstract

To identify cytoplasmic male sterility (CMS)-associated genes in tomato, we determined the genome sequences of mitochondria and chloroplasts in three CMS tomato lines derived from independent asymmetric cell fusions, their nuclear and cytoplasmic donors, and male fertile weedy cultivated tomato and wild relatives. The structures of the CMS mitochondrial genomes were highly divergent from those of the nuclear and cytoplasmic donors, and genes of the donors were mixed up in these genomes. On the other hand, the structures of CMS chloroplast genomes were moderately conserved across the donors, but CMS chloroplast genes were unexpectedly likely derived from the nuclear donors. Comparative analysis of the structures and contents of organelle genes and transcriptome analysis identified three genes that were uniquely present in the CMS lines, but not in the donor or fertile lines. RNA sequencing analysis indicated that these three genes transcriptionally expressed in anther, two of which were also expressed in pollen. They could be potential candidates for CMS-associated genes. This study suggests that organelle reorganization mechanisms after cell fusion events differ between mitochondria and chloroplasts, and provides insight into the development of new F1 hybrid breeding programs employing the CMS system in tomato.

## Introduction

Cytoplasmic male sterility (CMS) is broadly found in the kingdom of Plantae^1^. CMS plants cannot produce seeds by self-pollination due to a lack of male fertility; therefore, pollen from other plants is always required for these plants to produce seeds. CMS is caused by the incompatibility of interactions of genetic information between nuclei and organelles, especially mitochondria^1^. The genes in nuclei and organelles are called *restore of fertility* (*RF*) genes and CMS-associated genes, respectively. Therefore, CMS plants have been used as materials for studies of interactions between nuclear and cytoplasmic genes. Moreover, CMS is used in breeding programs to produce F1 hybrid seeds^1^, in which cytoplasmic and pollen donors are employed as maternal and paternal parents, respectively.

CMS plants can be artificially generated by recurrent backcrossing or transgenic approaches^2,3^, which leads to incompatibility between nuclei and organelles. A tomato CMS line, called CMS-pennellii, which possesses nuclei and cytoplasm from *Solanum pennellii* and *Solanum peruvianum*, respectively, has been developed by recurrent backcrossing^2^. A gene knockdown strategy is also used to develop CMS tomato lines, for which expression of a nuclear gene that regulates mitochondrial substoichiometric shifting has been suppressed^3^. In addition, other types of CMS tomato lines have been generated via asymmetric cell fusion between cultivated tomato lines, namely, *Solanum lycopersicum* as the nuclear donor and a wild potato relative, *Solanum acaule*, as the cytoplasmic donor^4^. Among CMS lines, MSA1 has been well-studied to reveal nucleus-organelle incompatibility^4^. A physical map of the mitochondrial genome of MSA1 indicates that this asymmetric cell fusion hybrid has a complex mitochondrial genome structure consisting of the parental genomes^5^. Transcripts of an open reading frame (ORF), *orf206*, of the hybrid mitochondrial genome are heterogeneously edited^6^. However, no candidates of CMS-associated genes have been identified in tomato.

Although CMS-associated gene sequences are not conserved across plant species, they have common features^7^. Most CMS-associated gene candidates usually possess transmembrane regions and chimeric structures, so-called fusion genes, of genes involved in respiration. Based on this information, CMS-associated genes have been identified in *Oryza sativa*^8,9^, *Helianthus annuus*^10^, and *Gossypium hirsutum*^11^. RNA-sequencing (RNA-Seq) based on next-generation sequencing technology has been employed to select candidates uniquely expressed in CMS lines of *Brassica juncea*^12^. Further functional studies are required to confirm that these candidates are involved in CMS. Introduction of *RF* genes into CMS lines would be a useful approach because CMS-associated genes can be downregulated in the presence of *RF* genes^7^. Another approach is to introduce CMS-associated genes into fertile lines to induce sterility^13^. More recently, it has become possible to alter or edit gene sequences of mitochondrial genomes with TALEN technology^14^. This technology has been used to disrupt CMS-associated genes in mitochondrial genomes and thereby generate *Arabidopsis thaliana*, *Oryza sativa*, and *Brassica napus* with CMS^14,15^.

In parallel with MSA1, as shown in Figure 1, two asymmetric cell fusions were developed between cultivated tomato lines *S. lycopersicum* (‘O’ and ‘P’) as nuclear donors and a wild potato relative, *S. acaule*, as the cytoplasmic donor^16^. The nuclear genome backgrounds of the three cell fusion lines including MSA1 were replaced with the genomes of cultivated tomato lines by a repeated backcrossing strategy. The resultant CMS lines are designated ‘CMS[MSA1]’, ‘CMS[O]’, and ‘CMS[P]’. Therefore, it may be possible to identify CMS-associated genes by comparative analysis of the genomes and transcriptomes of the CMS lines and their nuclear donors. In this study, we determined the sequences of the organelle genomes of the CMS lines and their donors. Subsequently, the genome sequences and gene expression patterns were compared to identify CMS-associated gene candidates. Furthermore, the results of this analysis may provide insights into the cytoplasmic genome features of asymmetric cell fusions.

**Figure 1.**
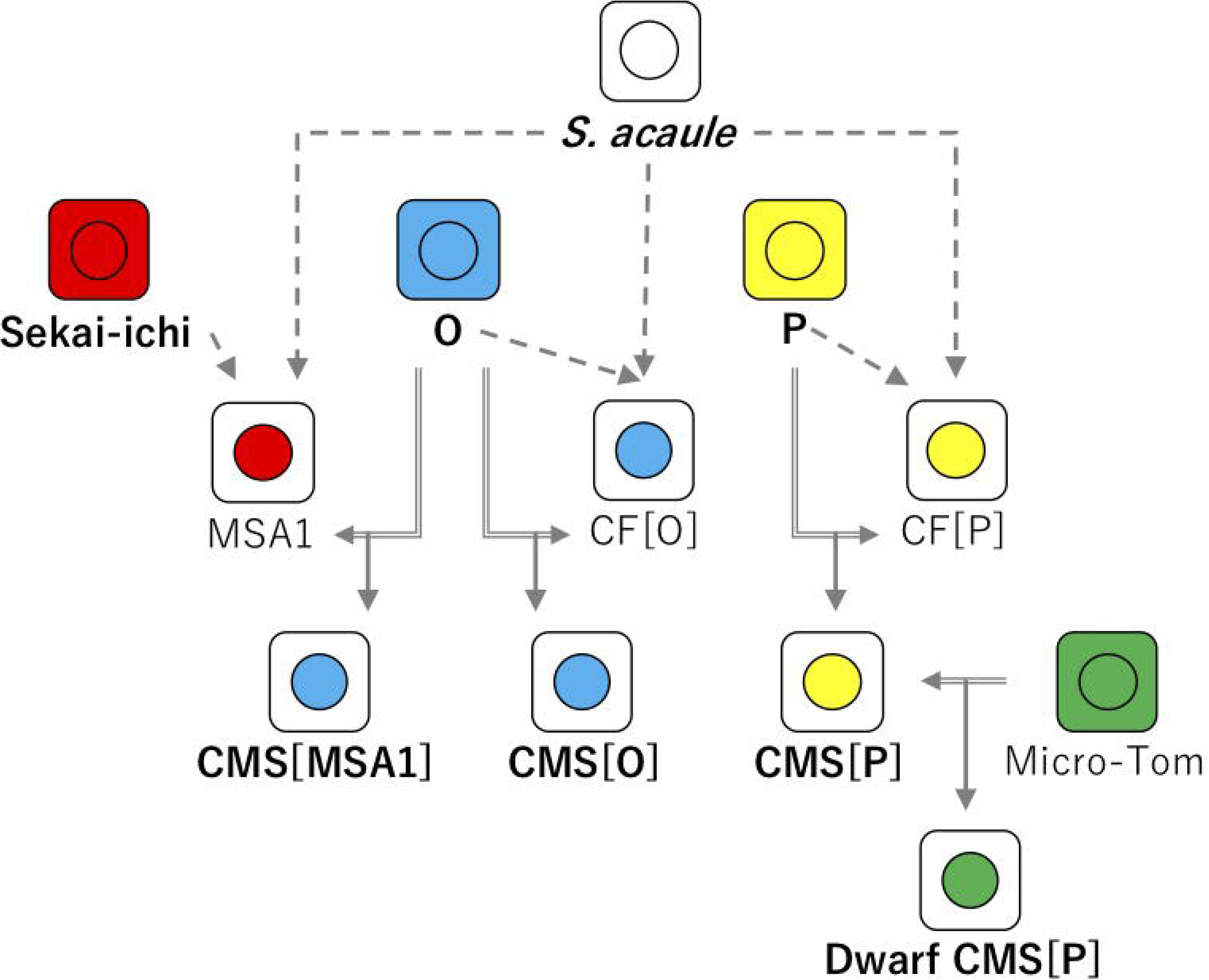
Pedigree of the CMS tomato lines. Squares and circles indicate cytoplasm and nuclei, respectively. Arrows with dashed lines and doubled lines indicate cell fusions and crossings, respectively.

## Results

### De novo assembly of chloroplast and mitochondrial genomes

A total of 10.5 Gb reads per sample were obtained from three CMS tomato lines (‘CMS[MSA1]’, ‘CMS[O]’, and ‘CMS[P]’), three nuclear donors (‘Sekai-ichi’, ‘O’, and ‘P’), and one cytoplasmic donor (*S. acaule*). Of them, 374 Mb (3.6%) and 566 Mb (5.4%) of reads per sample were aligned on publicly available sequences of mitochondrial and chloroplast genomes, respectively. The reads mapped on the two sets of reference sequences were separately assembled into contig sequences.

Mitochondrial genome sequences were constructed with reads mapped on the mitochondrial reference sequences (Table 1). The mitochondrial genomes of the nuclear donors ‘Sekai-ichi’, ‘O’, and ‘P’ were all constructed from only contigs with assembly sizes of 562.6 kb (*n* = 2, *n* represents contig numbers), 536.9 kb (*n* = 2), and 553.3 kb (*n* = 2), respectively. In *S. acaule*, 728.4 kb contigs (*n* = 7) for the mitochondrial genome were established. The assembly sizes were longer in the CMS lines than in the nuclear and cytoplasmic donors, specifically, they were 995.2 kb (*n* = 7) in ‘CMS[MSA1]’, 968.4 kb (*n* = 7) in ‘CMS[O]’, and 829.3 kb (*n* = 5) in ‘CMS[P]’. For chloroplast genomes, total sequence lengths of 389.2 kb (*n* = 2), 349.5 kb (*n* = 2), and 346.9 kb (*n* = 2) were constructed for ‘Sekai-ichi’, ‘O’, and ‘P’, respectively (Table 1). There were two contig sequences in each of the three nuclear donors. The assembly sizes were shorter in ‘CMS[MSA1]’ (296.6 kb, *n* = 1) and ‘CMS[O]’ (307.1 kb, *n* = 1) than in the nuclear donors, but longer in ‘CMS[P]’ (454.1 kb, *n* = 3).

**Table 1.**
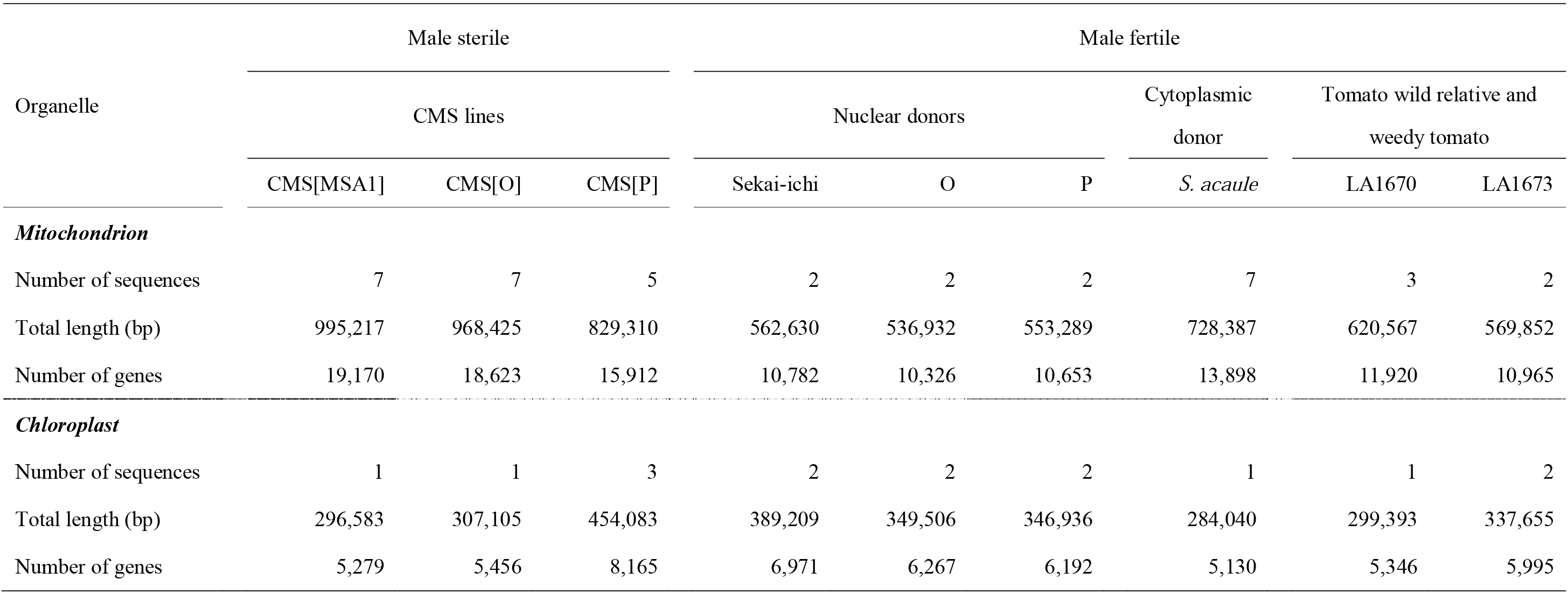
Assembly data of the organelle genomes.

Comparative genome analysis revealed that the mitochondrial genomes of the CMS lines consisted of highly fragmented, repeated, and duplicated sequences derived from both donors throughout the genome (Figure 2). On the other hand, the structures of the chloroplast genomes of the CMS lines were moderately conserved across the nuclear and cytoplasmic donors (Figure 2).

**Figure 2.**
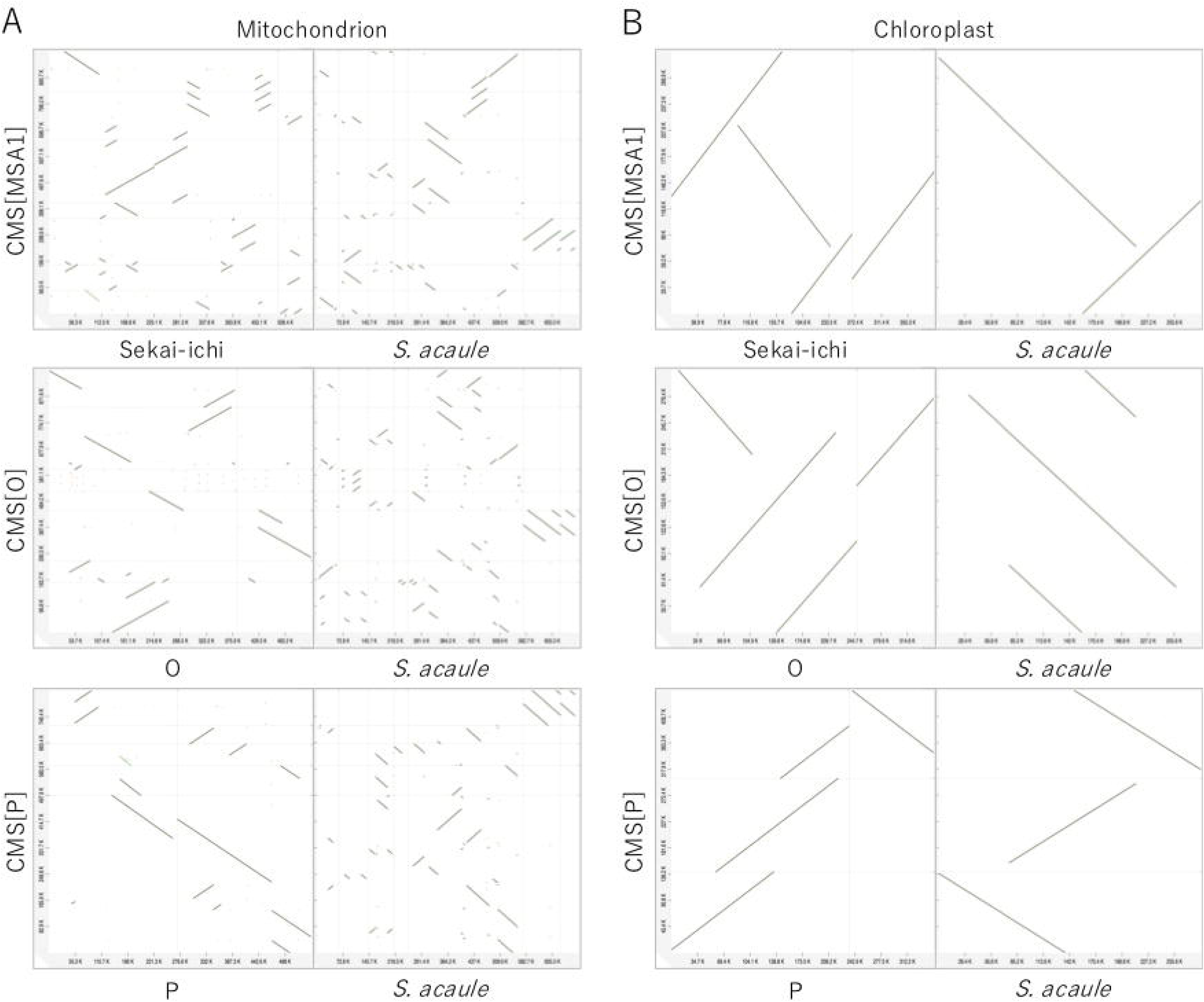
Comparative maps of the organelle genomes of the CMS tomato lines. Mitochondrial (A) and chloroplast (B) genomes of the three CMS lines, nuclear donors, and cytoplasmic donor. Dots indicate sequence similarity between the genome sequences.

In parallel, we determined the mitochondrial and chloroplast genome sequences of *Solanum pimpinellifolium* LA1670 and *S. lycopersicum* var. *cerasiforme* LA1673 (Table 1). Sequence reads were obtained from a public DNA database and processed as described above. Assembly sizes of the mitochondrial and chloroplast genomes were 620.6 kb (*n* = 3) and 299.4 kb (*n* = 1) for *S. pimpinellifolium* LA1670, respectively, and 569.9 kb (*n* = 2) and 337.7 kb (*n* = 2) for *S. lycopersicum* var. *cerasiforme* LA1673, respectively.

### Gene prediction from the organelle genomes

ORFs encoding ≥25 amino acids were extracted from the assembled sequences to predict potential genes. The number of potential genes predicted from the chloroplast genome assemblies ranged from 5,130 (*S. acaule*) to 8,165 (‘CMS[P]’) and the number of potential genes predicted from the mitochondrial sequences ranged from 10,326 (‘O’) to 19,170 (‘CMS[MSA1]’) (Table 1).

The ORFs were clustered to identify genes unique to and shared among the CMS lines, nuclear donors, and cytoplasmic donor (Figure 3). The ORFs in the CMS mitochondrial genomes consisted of four types of genes, namely, those unique to the CMS lines (Type 1: 9.4–11.9%), those shared with the nuclear donors only (Type 2: 14.1–17.0%), those shared with the cytoplasmic donor only (Type 3: 8.9–13.2%), and those shared with both the nuclear and cytoplasmic donors (Type 4: 61.8–64.1%). By contrast, the ORFs in the CMS chloroplast genomes mostly consisted of three types of genes, namely, those unique to the CMS lines (Type 1: 1.2–5.9%), those shared with the nuclear donors only (Type 2: 31.2–33.1%), and those shared with both the nuclear and cytoplasmic donors (Type 4: 62.9–65.7%). Few genes shared with the cytoplasmic donor only were found (Type 3: up to 0.1%).

**Figure 3.**
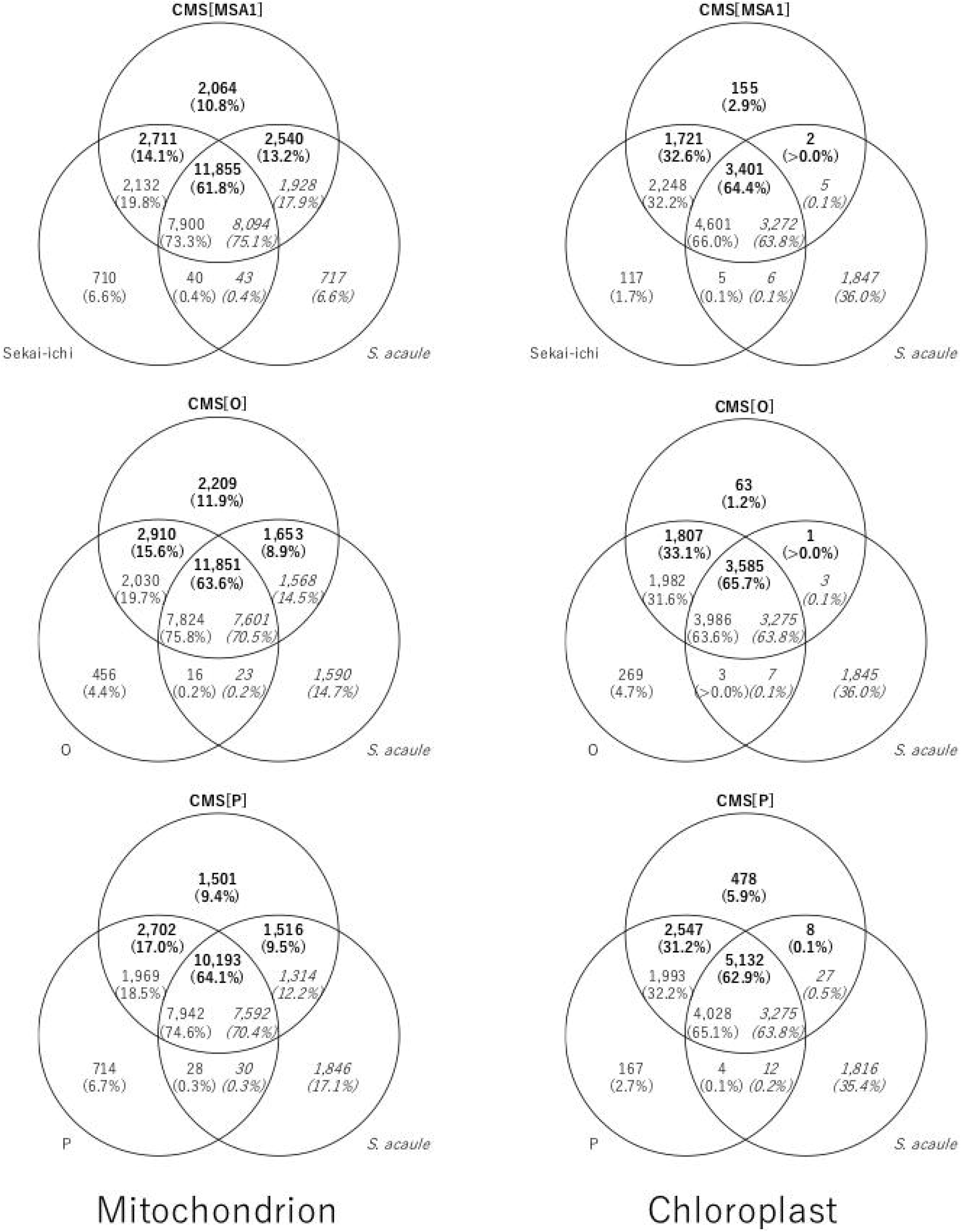
Organelle genes in the CMS lines, nuclear donors, and cytoplasmic donor. Numbers of genes unique to the CMS lines, nuclear donors, and cytoplasmic donor are indicated in bold, standard, and italic fonts, respectively. Percentages of genes are shown in parentheses.

The genome positions of the genes differed according to the gene type and organelle (Figure 4). Type 1 genes in mitochondria were distributed across the genome with some gaps. The positions of Type 2 genes were basically the same as those of Type 1 genes, while Type 3 genes were located in the gaps between Type 1 genes. Type 4 genes were also located in the gaps and at the ends of contig sequences. On the other hand, in chloroplast genomes, the positions of Type 1 and 2 genes overlapped and Type 4 genes were located at the ends of contigs.

**Figure 4.**
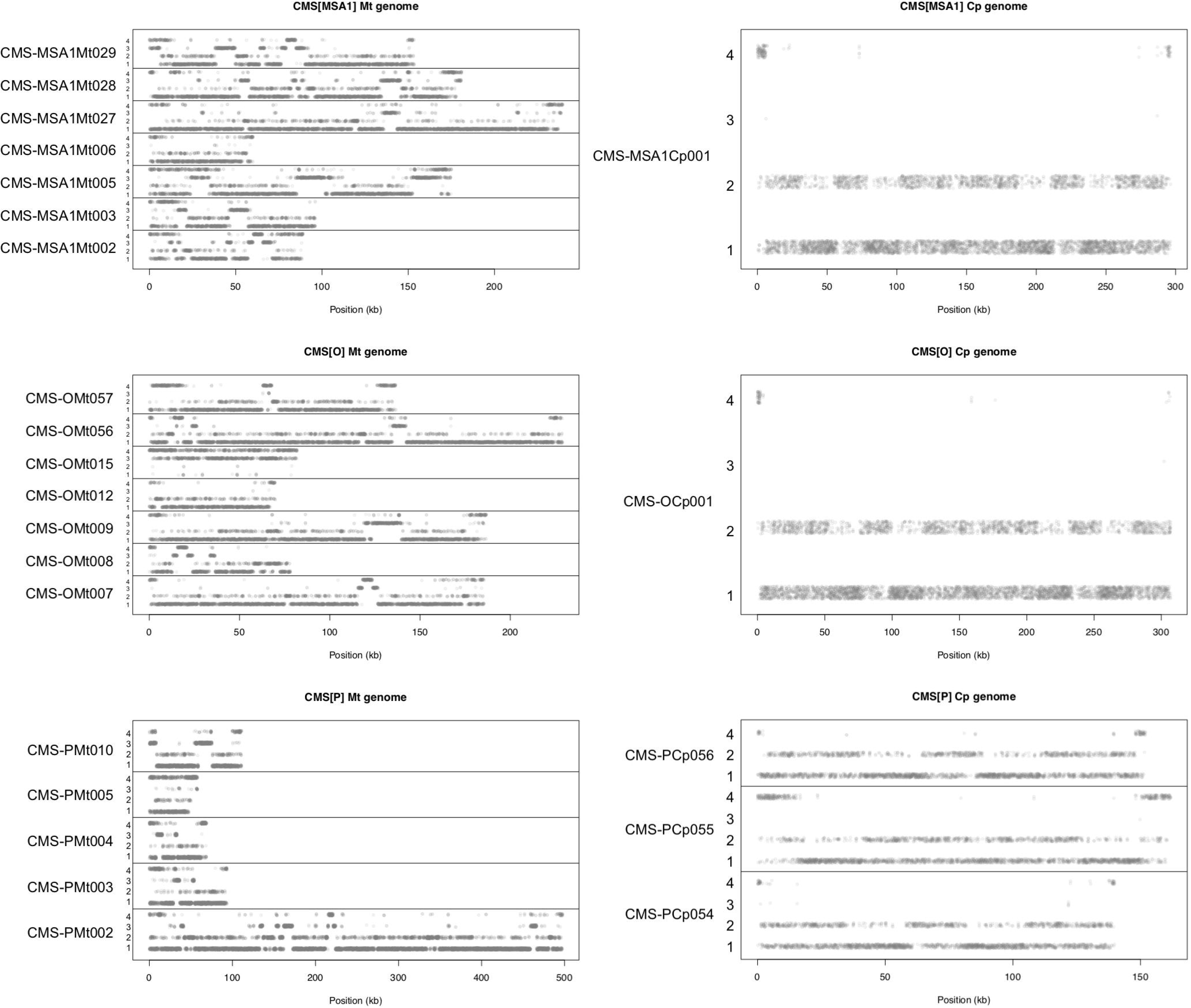
Distributions of CMS tomato genes across the organelle genomes. Dots indicate gene positions on contig sequences of the organelle genomes. Genes are grouped into the following four types: Type 1, genes unique to the CMS lines; Type 2, genes shared with the nuclear donors; Type 3, genes shared with the cytoplasmic donor; and Type 4, genes shared with both the nuclear and cytoplasmic donors.

### Screening of CMS-associated gene candidates

To identify candidates of CMS-associated genes in the mitochondrial genomes, we set the following four criteria: 1) amino acid length ≥70, 2) absent from male fertile lines, 3) present in all three CMS lines, and 4) expressed in anthers of the CMS lines. Among the predicted genes in the ‘CMS[P]’, ‘CMS[MSA1]’, and ‘ CMS[O]’ mitochondrial genomes, 831, 1,025, and 969 genes encoded ≥70 amino acids, respectively. The gene sequences from the CMS lines were compared with the mitochondrial genomes of the nuclear donors (‘Sekai-ichi’, ‘P’, and ‘O’) and *S. pimpinellifolium* LA1670, *S. lycopersicum* var. *cerasiforme* LA1673), *S. pennellii*, and *Nicotiana tabacum*. In total, 183, 272, and 140 genes were selected because they were absent from the nuclear donors and Solanaceae relatives, all of which possess male fertility. Furthermore, we selected 36, 41, and 33 genes commonly present in the CMS lines. The copy numbers of the genes varied. Finally, RNA-Seq reads were mapped on the mitochondrial genomes of the CMS lines. This analysis limited the number of CMS-associated gene candidates to four, including two identical sequences. The three genes were named *orf137* (two copies in the genome of each CMS line: CMS-PMt002g07240 and CMS-PMt005g13392), *orf193* (one copy: CMS-PMt002g06465), and *orf265* (one copy: CMS-PMt010g15739).

*De novo* transcriptome assembly was performed in parallel. RNA-Seq data were obtained from the anthers of ‘P’ and ‘CMS[P]’, and assembled into 62 and 43 transcript sequences, respectively, of which 37 ‘P’ and 18 ‘CMS[P]’ transcripts were predicted to have transmembrane domains. Of these sequences, eight were uniquely detected in ‘CMS[P]’. Two genes (STRG.32.1.p1 and STRG.39.1.p1) were identical to *orf137* and *orf265*.

Because two genes were commonly identified in both analyses, a total of nine genes were finally selected as candidates of CMS-associated genes (Table 2). Sequence similarity searches with the mitochondrial and chloroplast genomes indicated that two copies of the STRG.32.1.p1 (*orf137*) sequence (CMS-PMt002g07240 and CMS-PMt005g13392) were present in the mitochondrial genomes of the three CMS lines. A single copy sequence of *orf193* (CMS-PMt002g06465) and a single copy sequence of STRG.39.1.p1 (*orf265*, CMS-PMt010g15739) were found in the mitochondrial genomes of the three CMS lines in addition to that of *S. acaule*. The presence of the three genes in the CMS lines was validated by a PCR assay with the three CMS lines and six fertile lines. The remaining six genes were found in both the CMS and fertile lines. We selected three genes, *orf137*, *orf193*, and *orf265*, as highly potential candidates for CMS-associated genes due to their presence specifically in the CMS mitochondrial genomes and their expression in anthers.

**Table 2.**
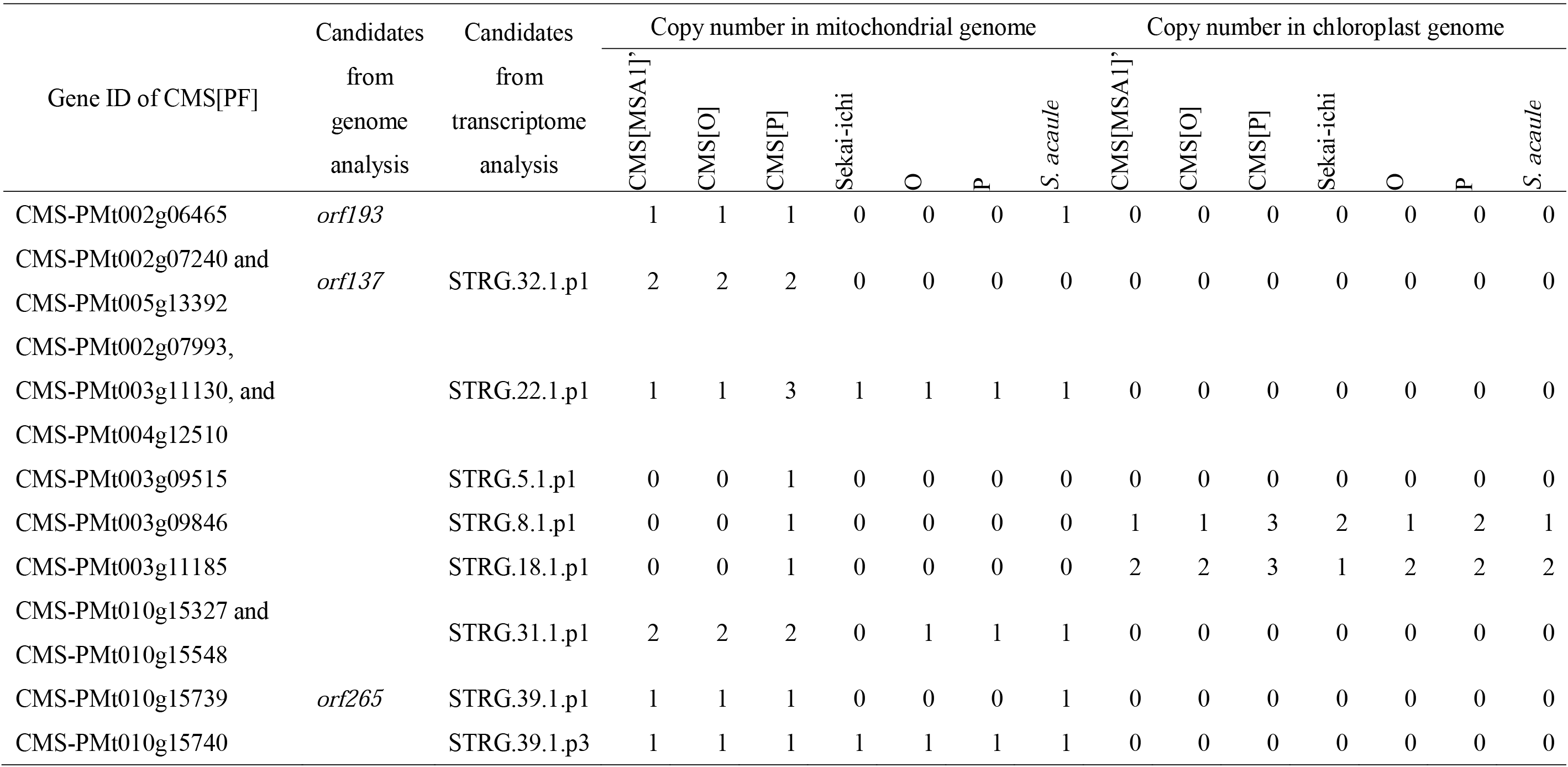
Copy numbers of CMS-associated gene candidates in the organelle genomes.

### Sequence similarity analysis of the candidate genes

The sequence similarity of the candidate genes including their flanking genome regions in the mitochondrial genome of ‘CMS[P]’ was investigated. A 3,045 bp genome sequence around *orf193* showed high sequence similarity to a 4,682 bp region of the tomato chloroplast genome sequences. The 3,045 bp sequence was split into three sequences containing 1,590, 488, and 1,007 bp (Figure 5A) with highly conserved boundary sequences (Figure 5B). In the 1,590 bp chloroplast genome sequence, a gene encoding *cytochrome f* was encoded; however, the corresponding sequence in the mitochondrial genome had a single base insertion causing a frame-shift mutation (Figure 5C). This mutation broke the ORF of the *cytochrome f* gene and generated two small ORFs, *orf116* and *orf193*.

**Figure 5.**
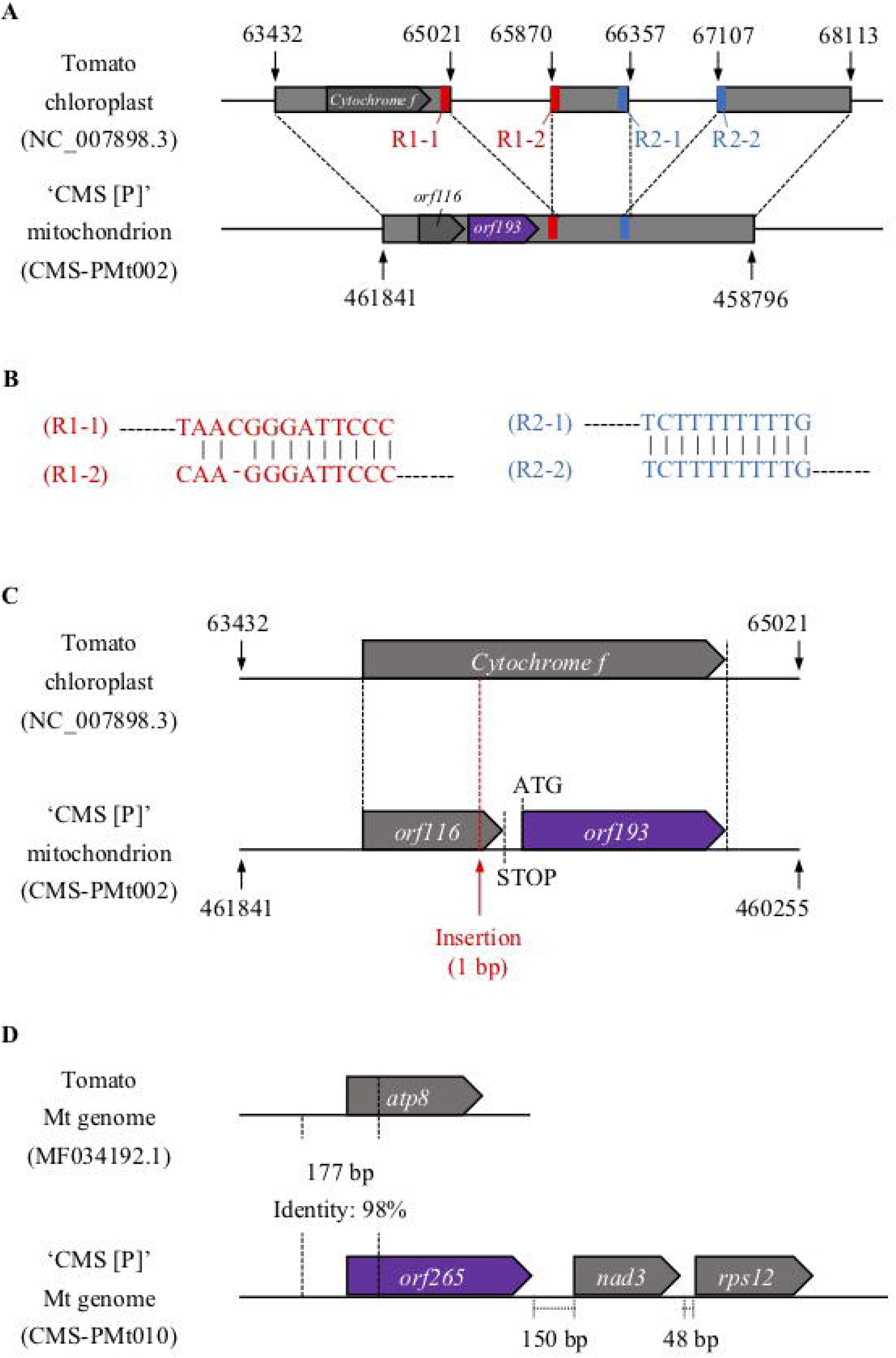
Structures of mitochondrial genes in ‘CMS[P]’. A. Genome structure of the *orf193* region. Homologous sequences between the two genomes are indicated by gray boxes. Highly conserved sequences at the borders are shown in red and blue. B. Sequence alignments of the borders. C. Details of the genome structure of the *orf193* region. A single nucleotide insertion causing a frame-shift mutation is indicated with a red arrow. D. Genome structure of the *orf265* region.

A portion of *orf265* and its upstream sequences (177 bp in total) showed high similarity to the *ATP synthase subunit 8* (*atp8*) gene encoded in the tomato mitochondrial genome (Figure 5D). The remaining sequences of *orf265* lacked similarity to reported sequences. *orf265* was located upstream of the *nad3* and *rps12* genes in the mitochondrial genome. No sequence similarity was observed for *orf137* and the flanking sequence.

### Expression analysis of the candidate genes

The expression patterns of the candidate genes, *orf137*, *orf193*, and *orf265*, were investigated by RT-PCR. First, we validated the results of the transcriptome analysis by detecting expression of the three genes in anthers of ‘CMS[P]’ and ‘CMS[MSA1]’ (Figure 6A). *orf265* was tandemly arrayed with *nad3* and *rps12*; therefore, we assumed that these three genes were co-transcribed as an operon. As expected, transcripts spanning the three genes were also detected (Figure 6A). Next, we analyzed gene expression in leaves, stems, roots, ovaries, and pollen in addition to anthers of Dwarf ‘CMS[P]’ which was a BC3 generation of ‘CMS[P]’ backcrossed with a tomato dwarf cultivar ‘Micro-Tom’. Expression of *orf137* and *orf265* was detected in all tested tissues, while that of *orf193* was observed in leaves, stems, roots, ovaries, and anthers (Figure 6B).

**Figure 6.**
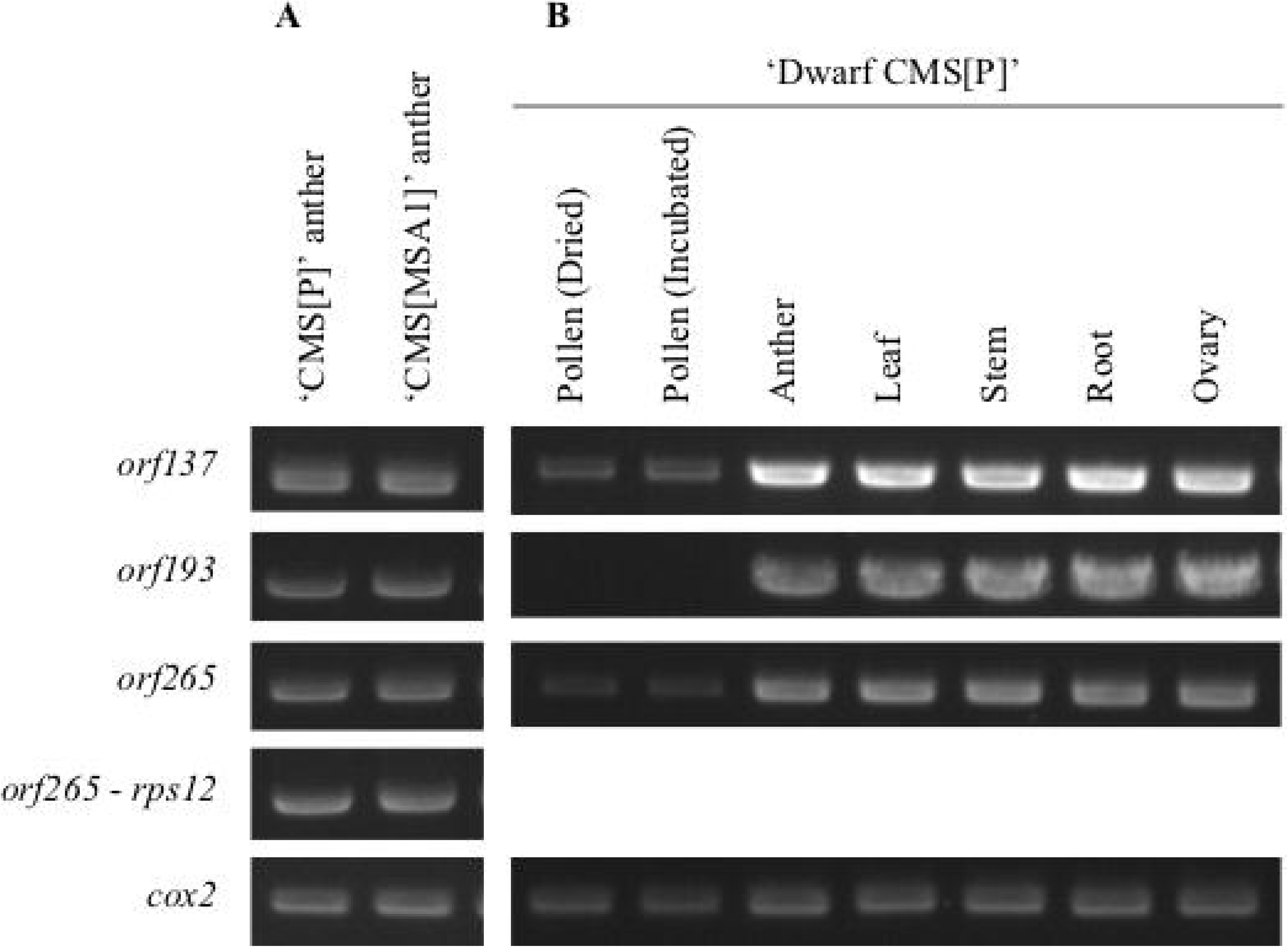
RT-PCR analysis of the CMS-associated gene candidates. Gene expression patterns in anthers of two CMS lines (A) and in seven samples of Dwarf ‘CMS[P]’ (B). *cox2* is a positive control.

## Discussion

We determined the mitochondrial and chloroplast genome sequences of CMS lines derived from asymmetric cell fusions and those of their nuclear and cytoplasmic donors (Table 1). Comparative analysis of the structures unexpectedly revealed that the cytoplasmic genome structures of the fusions were rearranged and divergent from those of the cytoplasmic donor (*S. acaule*) and nuclear donors (*S. lycopersicum*) (Figure 2). CMS-PMt003g09846 and CMS-PMt003g11185 were encoded in both the mitochondrial and chloroplast genomes (Table 2), suggesting that mitochondria and chloroplasts from the two donors were fused with each other and reorganized even though the cytoplasm of the nuclear donors was chemically inactivated to generate asymmetric cell fusions. Interestingly, the mitochondrial genomes of the CMS lines were larger than those of the donors, while the size of chloroplast genomes among the CMS lines were equivalent (Table 1). In addition, the structures of the mitochondrial genomes were divergent, while those of the chloroplast genomes were rather conserved (Figure 2). Gene clustering analysis suggested that both the cytoplasmic and nuclear donors contributed to form mitochondria in the CMS lines (Figure 3). Furthermore, the structures of the CMS mitochondrial genomes contained patches of the two genomes of the donors (Figure 4). These results suggest that the mitochondrial genomes of both donors were highly fragmented at the time of asymmetric cell fusion and reorganized to form a new mitochondrial genome^5^. This is completely different from our expectation that genomes of the cytoplasmic donors should be present in CMS lines derived from asymmetric cell fusions. More interestingly, chloroplasts of the CMS lines consisted only of genes from nuclear donors, not from the cytoplasmic donor. This unexpected finding has been frequently made in tomato^17^, tobacco^18^, and *Brassica*^19^. We speculate that interactions of genetic information between nuclei and organelles might be strict with chloroplasts rather than with mitochondria. The genome and/or organelle reorganization mechanisms after cell fusions might differ between mitochondria and chloroplasts.

Based on genome and transcriptome analyses, nine genes encoded in the mitochondrial genome of ‘CMS[P]’ were selected as candidate CMS-associated genes (Table 2). Among them, three genes (*orf193*, STRG.32.1.p1 = *orf137* and STRG.39.1.p1 = *orf265*) were uniquely present in the genomes of the CMS lines and expressed in their anthers (Table 2 and Figure 6). STRG.32.1.p1 (*orf137*) showed sequence similarity with the CMS-associated protein encoding *cytochrome c subunit 1* (Figure 5). STRG.39.1.p1 (*orf265*) was similar to *ATP synthase subunit 8* at the N-terminus, but lacked similarity in the remaining regions (Figure 5). CMS-associated genes are generally involved in cellular respiration producing energy to generate pollen^20^, and this is true of both these genes. In many cases, fusion genes have been reported to be CMS-associated genes, e.g., *orf307* in *Oryza sativa*^21^ and *orf72* in *Brassica oleracea*^22^, and to produce cytotoxic proteins, which lead to male sterility. Knockout mutagenesis with mitoTALENs^14,15^ targeting the candidate genes would be useful to identify CMS-associated genes and to generate CMS lines from normal tomato cultivars.

CMS lines are powerful tools to produce F1 hybrid seeds in breeding programs^1^. However, in cereals and fruits including tomato, the *RF* genes are essential for F1 plants to set seeds and bear fruits. Restorer genes for CMS lines have been identified in wild tomato relatives, e.g., *S. pimpinellifolium* LA1670 and *S. lycopersicum* var. *cerasiforme* LA1673^16^. Recently, we published the genome sequence data of these two wild relatives^23^. We expect *RF* genes for CMS lines to be discovered soon based on this information, although no candidate genes or genetic loci have been reported. Once CMS-associated genes and restorer genes are identified, tomato F1 hybrid seeds can be produced by employing insect pollinators instead of the currently used hand-pollination systems. We propose that CMS-based F1 hybrid breeding programs with insect pollinators can be implemented in tomato breeding programs to reduce the costs of F1 seed production in the future.

## Materials and methods

### Plant materials

Three tomato CMS lines (‘CMS[MSA1]’, ‘CMS[O]’, and ‘CMS[P]’), three cultivated tomato lines (*S. lycopersicum* ‘Sekai-ichi’, ‘O’, and ‘P’), and one potato wild relative (*S. acaule*) were used (Figure 1). ‘ CMS[MSA1]’ was developed by repeated backcrossing using ‘O’ as a recurrent parent and a male-sterile tomato, MSA1, as a cytoplasmic donor. MSA1 is an asymmetric cell fusion between the tomato cultivar Sekai-ichi (as the nuclear donor) and the potato wild relative *S. acaule* (as the cytoplasmic donor)^4^. ‘CMS[O]’ was a progeny in repeated backcrossing using ‘O’ as the paternal parent and an asymmetric cell fusion between ‘O’ (as the nuclear donor) and *S. acaule* (as the cytoplasmic donor). ‘CMS[P]’ was also a progeny in backcrossing using ‘P’ as the paternal parent and an asymmetric cell fusion between ‘P’ (as the nuclear donor) and *S. acaule* (as the cytoplasmic donor). Dwarf ‘CMS[P]’ was developed from ‘CMS[P]’ by backcrossing with *S. lycopersicum* ‘Micro-Tom’ (TOMJPF0001), which is a miniature dwarf cultivar^24^. The putative nuclear and cytoplasmic genomes of the materials are shown in Figure 1.

### Genome sequence analysis

Total genomic DNA was extracted from young leaves of the six tomato lines (‘CMS[MSA1]’, ‘CMS[O]’, ‘CMS[P]’, ‘Sekai-ichi’, ‘O’, and ‘P’) and *S. acaule* with a Maxwell 16 Instrument and Maxwell 16 Tissue DNA Purification Kits (Promega, Madison, WI, USA). SMRT sequence libraries were constructed with an SMRTbell Express Template Prep Kit (PacBio, Menlo Park, CA, USA) and used for sequencing on a PacBio Sequel system (PacBio). Genome sequence data for S. *pimpinellifolium* LA1670 and *S. lycopersicum* var. *cerasiforme* LA1673 were obtained from a public DNA database (DRA accession numbers DRX231405 and DRX231409)^23^.

### Genome assembly and gene prediction

Sequence reads were mapped on reference genome sequences for mitochondria (GenBank accession numbers MF034192, MF034193, NC_035964, and MF98995–MF989957) or chloroplasts (NC_007898) with Organelle_PBA^25^. Reads mapped on the reference sequences were assembled into contig sequences with Canu^26^. Potential sequence errors in the contig sequences were corrected twice with the sequence reads by Arrow (PacBio). The corrected contig sequences were aligned back to the reference sequences with Nucmer^27^ to select highly confident organelle genomes. ORFs (≥75 bases) in the organelle genomes were selected as potential genes with ORFfinder (https://www.ncbi.nlm.nih.gov/orffinder). The ORF sequences were clustered with CD-HIT^28^. Transmembrane domains in the gene sequences were predicted by TMHMM^29^. Sequence similarity searches with the mitochondrial genomes of *S. pennellii* (NC_035964) and *N. tabacum* (NC_006581) were performed by BLAST^30^ with a threshold E-value of 1e-50.

### RNA expression analysis

Total RNA was extracted from the anthers of P and ‘CMS[P]’ with an RNeasy Plant Mini Kit (QIAGEN, Hilden, Germany). RNA was treated with RNase-free DNase (QIAGEN) and used for sequence library preparation with a TruSeq Stranded mRNA Library Prep Kit (Illumina, San Diego, CA, USA). The resultant libraries were sequenced on NextSeq500 (Illumina) in paired-end, 151 bp mode. After trimming adaptors and low-quality reads by Trim_galore (https://github.com/FelixKrueger/TrimGalore) with option -q 30 --length 100 followed by fastp^31^ with option -l 100, transcriptomes were *de novo* assembled by the HiSat2-Stringtie pipeline^32^ and putative ORFs were searched for annotation by BLASTP^30^ against the SWISS-PROT database^33^.

To validate the RNA-Seq results, RT-PCR was performed. In total, 800 ng of total RNA isolated from anthers of the CMS lines or seedlings of *S. acaule* was converted into cDNA with ReverTra Ace (TOYOBO, Osaka, Japan) using a random primer (TAKARA BIO, Kusatsu, Japan). cDNA diluted 10-fold with water was used as a template for PCR. The PCR mixture (10 μL) contained 0.5 μL cDNA, 0.3 μM primers (Table 3), 2× PCR buffer (TOYOBO), 400 μM dNTPs, and 1 U DNA polymerase (KOD FX Neo, TOYOBO). The thermal cycling conditions were as follows: initial denaturation at 94°C for 3 min; 35 cycles of denaturation at 98°C for 15 s, annealing at 60◻ for 30 s, and extension at 68°C for 60 s; and a final extension at 68°C for 3 min. PCR products were separated by electrophoresis in a 1% agarose gel with TAE buffer. Gels were stained with Midori Green Advance (NIPPON Genetics, Tokyo, Japan) to detect DNA bands under ultraviolet illumination.

**Table 3.**
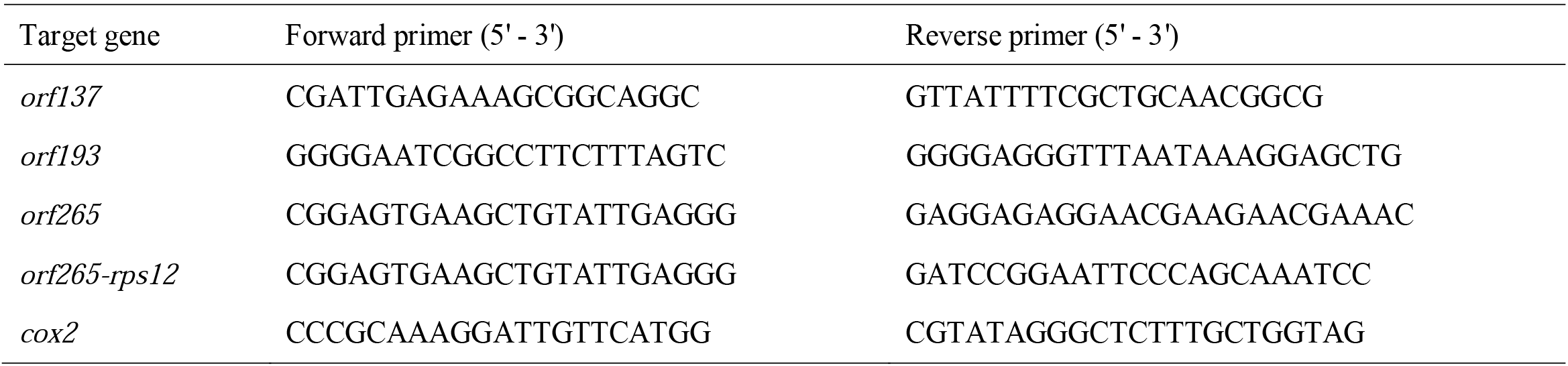
Oligonucleotide sequences of PCR primers

## Data availability

The DDBJ accession numbers of the assembled sequences are LC613090-LC613141. Genome information is available at KaTomicsDB (http://www.kazusa.or.jp/tomato).

## Acknowledgments

We are grateful to Prof. Kazuo Watanabe (University of Tsukuba, Japan) for providing the DNA material of *S. acaule*. This work was supported by the Kazusa DNA Research Institute Foundation and the Project of the NARO Bio-oriented Technology Research Advancement Institution (Research Program on Development of Innovative Technology, Grant number: 30010A). Seeds of Micro-Tom (TOMJPF0001) was provided from National BioResource Project Tomato (NBRP tomato).

## Contributions

TA and KS conceived and coordinated the project. YM established the plant materials. KK, IH, and KS collected the data. KK, IH, TA, and KS analyzed and interpreted the data. KK and KS wrote the manuscript with contributions from TA. All authors read and approved the final manuscript.

